# HaloMPNN: retraining ProteinMPNN on halophilic proteomes for salt-tolerant enzyme design

**DOI:** 10.64898/2026.08.02.742362

**Authors:** Alyssa Lu Lee, Austin Seamann, Gwendolyn Chung, Clairie Zhao, Rohan Maddamsetti, Sagar D. Khare

**Author notes:** These authors contributed equally to this work (Alyssa Lu Lee, Austin Seamann).

## Abstract

Machine learning-guided protein sequence redesign is now routinely used to optimize multiple properties relevant for protein engineering, most prominently thermostability and recombinant expression levels. Salt tolerance is a valuable property for ”blue-biotechnology”- enabled biomanufacturing, yet no generative computational method exists to redesign proteins for increased salt tolerance. We hypothesized that a training dataset heavily biased toward salt-adapted proteomes would yield a model capable of designing proteins with halophilic properties. To test this, we retrained the sequence redesign model ProteinMPNN on proteins from “salt-in” extreme halophiles such as *Haloarcula marismortui*, a Dead Sea archaeon that grows optimally near 3–4 M NaCl, roughly six times the salinity of seawater, and accumulates molar concentrations of salts in its cytoplasm. Our model, HaloMPNN, redesigns non-halophilic proteins so that their properties shift towards those of natural halophilic proteins: lower predicted isoelectric point, greater surface acidity, and reduced surface and core hydrophobicity. Redesigning a broad range of non-halophilic proteins with SolubleMPNN, ProteinMPNN, and HyperMPNN shows that this shift is specific to HaloMPNN rather than a generic consequence of sequence redesign. HaloMPNN therefore offers both a route to designing candidate salt-tolerant enzymes and a means of identifying the characteristics that underlie halophilic adaptation.

## INTRODUCTION

Proteins from extremophiles remain folded and active under conditions that denature typical mesophilic proteins. Inverse folding models, structure-based neural networks that predict aminoacid sequences compatible with a given protein backbone, can be trained on such proteins to serve two complementary purposes. First, they are instruments for understanding adaptation: because a model such as ProteinMPNN^1^ reproduces the sequence distribution of its training set, retraining it on a given extremophile class concentrates the structural and compositional signatures of that class in its designs and, through the identification of accrued substitutions, generates hypotheses about which residues underlie extremophilic adaptation. Second, a retrained inverse folding model may offer a general route for designing enzymes that function at high temperature, extreme pH, or high salinity. This retrain-on-phenotype strategy has been demonstrated experimentally for solubility ^2^ and thermostability^3^, and proposed computationally for alkaline adaptation^4^, but how generally it applies across protein properties, and whether its designs function as intended, remains an open question. A single retrained model, however, only demonstrates that a training distribution can be reproduced. Comparing models trained on different phenotypes against the same target proteins is more informative, because it separates shifts specific to the intended adaptation from those that a generic inverse folding model would produce. This comparison, therefore, tests whether the model has captured adaptation-relevant structure-to-sequence relationships rather than a generic compositional bias.

Among extreme environments, high salinity is of particular interest for biomanufacturing^5,6^. Enzymes that stay soluble and active at molar salt concentrations enable processes that run in seawater or brine rather than fresh water, tolerate the high ionic strength of many industrial feedstocks, and resist microbial contamination under conditions in which most organisms cannot grow^6,7^. Such capabilities are central to “blue biotechnology”, the use of marine and other aquatic bioresources for sustainable production^8^. Halophilic microorganisms have already solved the underlying biophysical problem, since the cytoplasmic proteins of salt-in halophiles function at intracellular salt concentrations that would aggregate or unfold typical mesophilic proteins^9^. Learning the sequence features responsible for this adaptation and transferring them to proteins of interest would therefore provide a general route to salt-tolerant biocatalysts. However, no generative computational method for designing salt-tolerant sequences currently exists.

Previous work characterizing natural halophilic proteins has identified several properties that distinguish them from non-halophilic homologs. The most prominent is a high percentage of negatively charged residues on the surface, which organize hydrated salt ions^10,11^, together with a reduction in solvent-exposed hydrophobic surface area^12^ and in buried hydrophobic contact area^13^. Rational engineering guided by these principles has increased the salt tolerance of individual enzymes while retaining function: substituting surface residues of a mesohalophilic carbonic anhydrase with acidic ones, a strategy rationalized by molecular dynamics simulations, produced an extreme halotolerant biocatalyst^14^. However, such rational design is not scalable and lacks the inferential power of deep learning, and it remains disputed whether increased surface negative charge or decreased hydrophobicity is individually necessary or sufficient for salt tolerance. Deep learning models can identify the subtle patterns associated with halotolerance, but so far they have been applied only to the classification, not the generation, of salt-tolerant proteins^15^, which motivates a data-driven approach that learns the relevant features jointly from natural halophilic proteins.

Here we test whether the retrain-on-phenotype strategy extends to halophilic adaptation. We assembled a dataset of 40,000 high-quality predicted structures drawn from extreme-halophile organisms in the HaloDom database, and retrained ProteinMPNN on it to produce HaloMPNN. To define the target signature, we first characterized the differences between halophilic and non-halophilic proteins across 1,433 paired homologs, confirming the expected shifts toward lower predicted isoelectric point, greater surface acidity, and reduced surface and core hydrophobicity. We then redesigned the non-halophilic members of these pairs with HaloMPNN and found that the redesigns recover this halophilic signature. Comparing HaloMPNN against ProteinMPNN, SolubleMPNN, and HyperMPNN on the same targets, we find that HaloMPNN reproduces the natural halophilic shift most closely, whereas the other MPNN models do not. The halophilic shift is therefore specific to HaloMPNN’s training set. More broadly, HaloMPNN is a step towards a general computational approach for increasing the salt tolerance of proteins, with broad applications in industrial biocatalysis and the potential to reveal the subtle molecular adaptations that underlie this phenotype.

## RESULTS

### Dataset curation and evaluation

To assemble a dataset of halophilic proteins to train HaloMPNN, we downloaded all protein sequences from organisms labeled “extreme” halotolerant in the HaloDom database^16^ and their corresponding predicted structures from AlphaFoldDB^17^, filtered for quality and length (Figure 1). Predicted membrane proteins and proteins in the evaluation dataset were excluded, and the remaining sequences were downsampled, clustered, and split (Methods). Proteins from extreme halophiles had substantially lower predicted isoelectric points than those from moderately or slightly halotolerant organisms (Figure S1), confirming that the training set was enriched for the acidic proteomes characteristic of salt-in adaptation. Trained on this dataset with the same architecture as Protein- MPNN^1^ (Figure 1), HaloMPNN converged over 1,000 epochs to a native sequence recovery of 52% on the training set and 50% on held-out validation sequences (Figure S2). The small gap between the two indicates that it learned generalizable halophilic sequence preferences rather than memorizing the training data.

**Figure 1:**
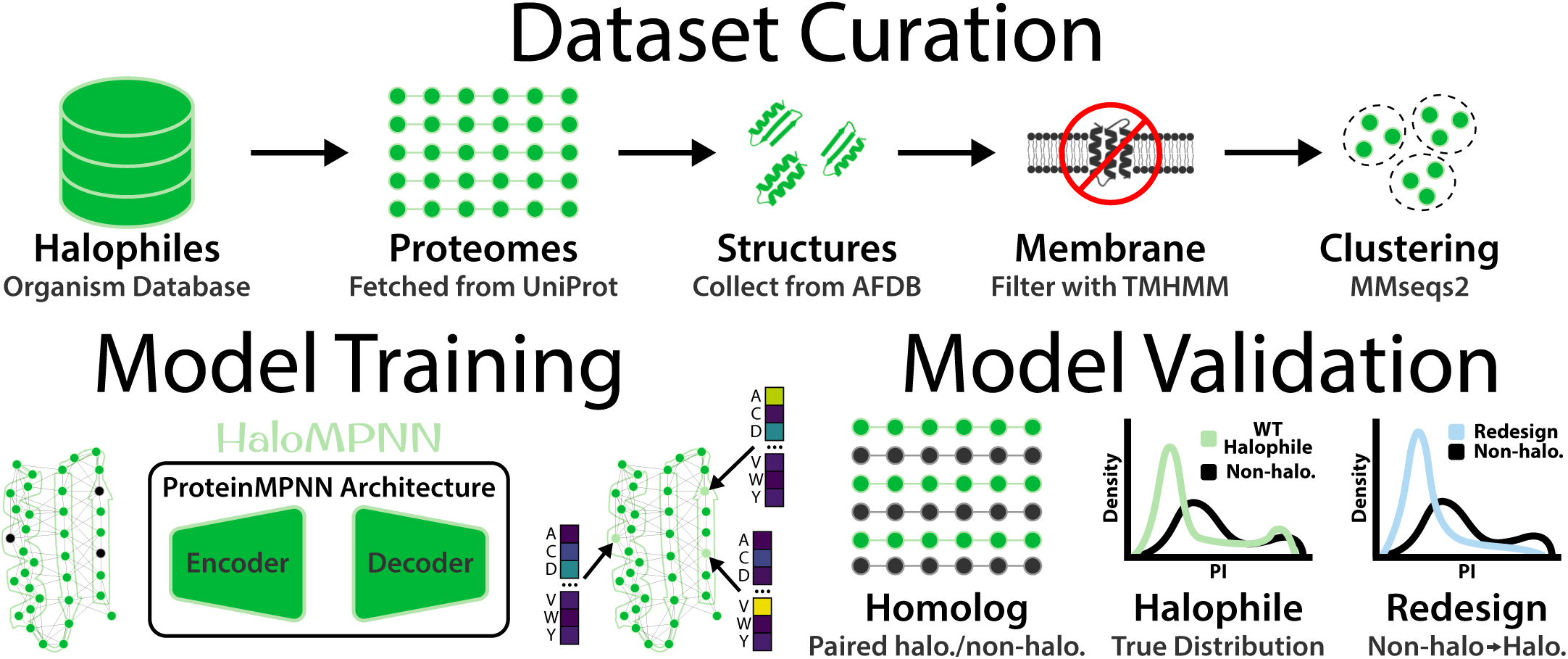
Overview of the HaloMPNN workflow. Halophilic protein sequences from organisms labeled “extreme” halotolerant in the HaloDom database ^16^, together with their AlphaFold-predicted structures ^17^, were filtered for quality and length, stripped of predicted membrane proteins and of the held-out evaluation set, then downsampled, clustered, and split into training, validation, and test sets (Methods). HaloMPNN was trained on these structures using the ProteinMPNN architecture ^1^. The model was evaluated on a curated set of paired halophilic and non-halophilic homologs by redesigning the non-halophilic members and comparing their properties with those of the natural halophilic proteins (Methods).

To determine which properties separate halophilic from non-halophilic proteins in nature, we reproduced and extended a result from Paul et al. ^10^ (Methods): sequence search identified 1,433 halophile/non-halophile homolog pairs across four pairs of species (Table S1). For each protein we computed the predicted isoelectric point and, from its predicted structure, the percentage of surface residues that are negatively charged, positively charged, or large hydrophobic, together with the large hydrophobic percentage of the buried core (Methods). We then redesigned the 1,433 non-halophilic proteins with HaloMPNN, holding fixed the positions conserved at *≥*50% in each protein’s multiple sequence alignment, and compared them with their natural halophilic counterparts and, for reference, with redesigns by ProteinMPNN^1^, SolubleMPNN^2^ (trained with membrane proteins excluded), and HyperMPNN^3^ (trained on hyperthermophilic proteins) using an identical list of designable positions. To assess generality, we repeated the redesign and analysis on 4,229 proteins from the *Escherichia coli* proteome with HaloMPNN, ProteinMPNN, and SolubleMPNN.

### Natural halophilic proteins have properties distinct from non-halophilic homologs

To investigate the molecular basis of halophilic adaptation, we compared each property between the halophilic and non-halophilic members of the 1,433 homolog pairs. The two groups separated clearly, and in the directions reported for salt-adapted proteins in the existing literature (Figure 2A; Tables 1 and 2). As expected, the halophilic proteins in our dataset have a lower predicted isoelectric point than their non-halophilic homologs. The bias towards negative surface charge is also reflected in our analysis. Compared to their non-halophilic homologs, halophilic proteins have a higher percentage of negatively charged surface residues and a lower percentage of positively charged surface residues. We also observe a lower percentage of large hydrophobic residues on the surface and in the core. (All five metrics differ at *q <* 10*^−^*^3^; Table S8.)

**Figure 2:**
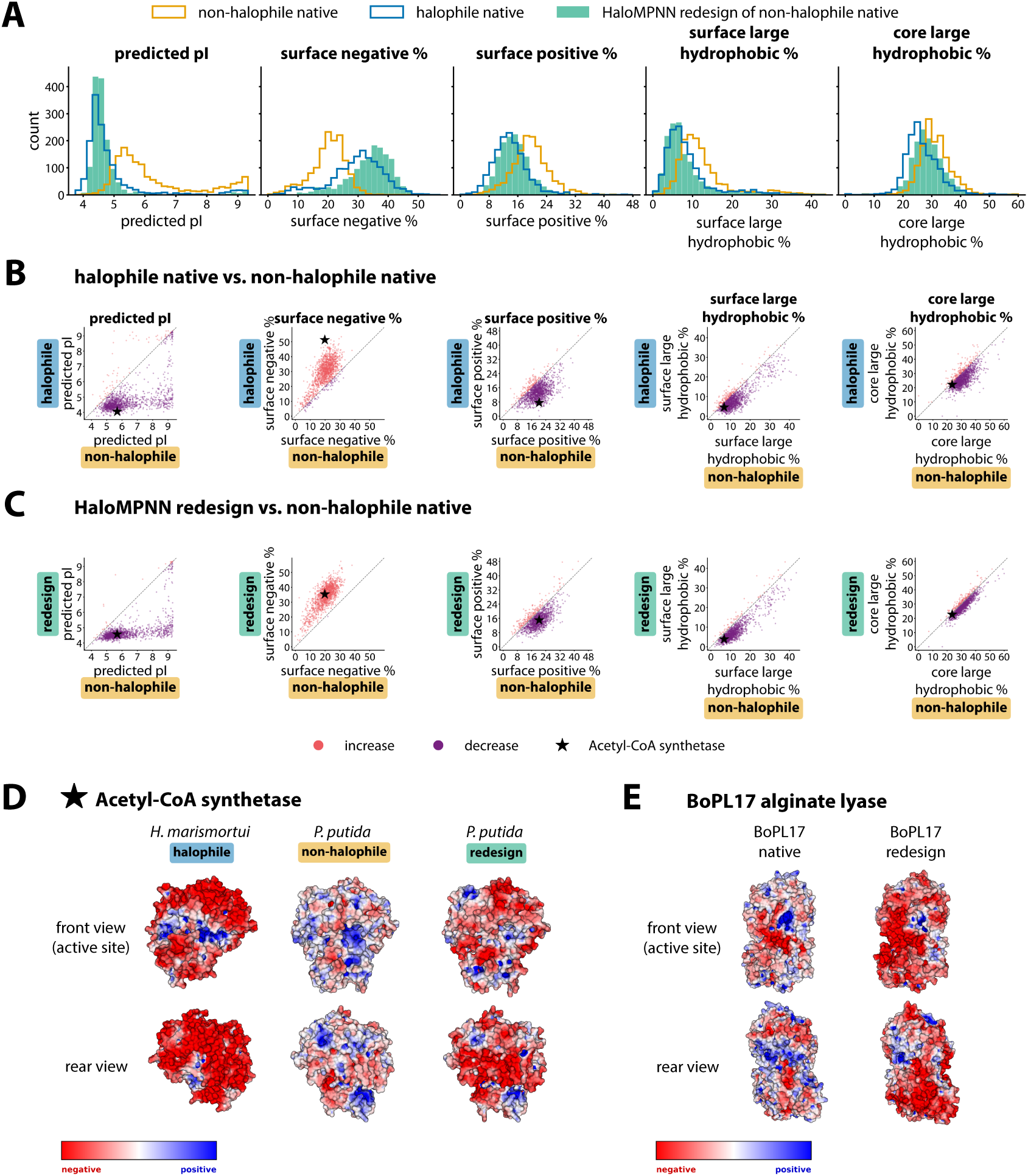
HaloMPNN redesigns non-halophilic proteins to match distinctive molecular properties of halophilic proteins. (A,B,C) Halophilic proteins show lower predicted isoelectric point, higher % negatively charged surface residues, lower % positively charged surface residues, lower % large hydrophobic surface residues, and lower % large hydrophobic core residues compared to their nonhalophilic homologs (*q <* 10*^−^*^3^ for all metrics; Table 1, Table S8). HaloMPNN redesigns of non-halophilic homologs show lower predicted isoelectric point, higher % negatively charged surface residues, lower % positively charged surface residues, lower % large hydrophobic surface residues, and lower % large hydrophobic core residues compared to pre-redesign (*q <* 10*^−^*^3^ for all metrics; Table 1, Table S8). Proteins shown are 1,433 pairs of homologs from halophilic and non-halophilic species identified in Table S1, plus HaloMPNN redesigns of the non-halophilic proteins. (D) Illustrative example: Acetyl-CoA synthetase from halophile *Haloarcula marismortui* has a higher density of negative charge on the surface compared to the same enzyme from non-halophile *Pseudomonas putida*. HaloMPNN redesign of acetyl-CoA synthetase from non-halophile *Pseudomonas putida* also has a high density of negative charge. Structures shown: AlphaFold predicted structure of Q5UXS3 (*Haloarcula marismortui*), AlphaFold predicted structure of Q88EH6 (*Pseudomonas putida*). (E) HaloMPNN redesign of alginate lyase BoPL17 from non- halophile *Bacteroides ovatus* has a higher density of negative charge on the surface compared to before redesign.

**Table 1:** Median predicted pI, surface negative %, surface positive %, surface large hydrophobic %, and core large hydrophobic % for 1,433 halophilic proteins and 1,433 non-halophilic homologs. Halophilic and non-halophilic species are listed in Table S1.

|  | predicted pI | surface negative % | surface positive % | surface large hydrophobic % | core large hydrophobic % |
| --- | --- | --- | --- | --- | --- |
| median (non-halophile) | 5.71 | 20.74 | 19.39 | 10.68 | 30.65 |
| median (halophile) | 4.53 | 30.63 | 13.46 | 7.04 | 25.70 |

**Table 2:**
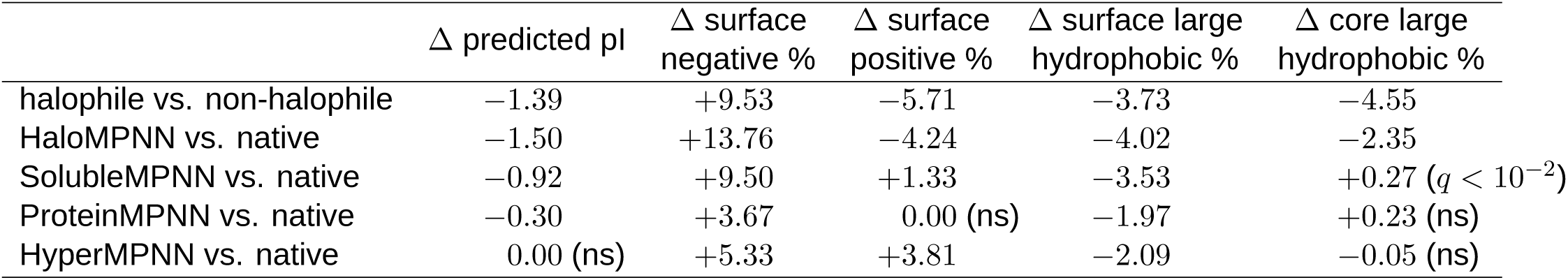
Mean change in predicted pI, surface negative %, surface positive %, surface large hydrophobic %, and core large hydrophobic %. The first row compares natural halophilic to non-halophilic homologs; the remaining rows compare each non-halophilic protein before and after redesign with the indicated model. All mean changes in protein properties are highly significant by a Benjamini–Hochberg-corrected sign test at *q <* 10*^−^*^3^ unless otherwise noted (ns, not significant) (Table S8).

Direct comparison of each halophilic protein with its non-halophilic homolog shows the same trends (Figure 2B). For almost every homolog pair, the predicted isoelectric point of the halophilic protein is lower than its non-halophilic homolog. Likewise, nearly every halophilic protein shown has a higher surface negative percentage, lower surface positive percentage, lower surface large hydrophobic percentage, and lower core large hydrophobic percentage compared to its non-halophilic homolog.

### HaloMPNN redesign of non-halophilic proteins approximates the properties of natural halophilic proteins

Redesigning the 1,433 non-halophilic proteins with HaloMPNN shifted their properties toward those of natural halophilic proteins across the entire set (Figure 2A; Table 2). Predicted isoelectric point fell, the fraction of negatively charged surface residues rose, and the fractions of positively charged and large hydrophobic surface residues both dropped, as did the large hydrophobic content of the core. All five metrics moved in the halophilic direction at *q <* 10*^−^*^3^ (Table S8).

These changes were consistent across the dataset rather than driven by a subset of proteins. In a paired before-and-after comparison, the predicted isoelectric point decreased for almost every one of the 1,433 targets, and the surface-negative, surface-positive, surface-hydrophobic, and core-hydrophobic percentages each moved in the halophilic direction for a comparable majority of proteins (Figure 2C).

Two enzymes illustrate the effect on individual proteins. Acetyl-CoA synthetase, selected as an illustrative example (Methods), differs markedly between the halophile *Haloarcula marismortui* and the non-halophile *Pseudomonas putida*, with the halophilic enzyme carrying far more negative charge on its surface (Figure 2D). Redesigning the *P. putida* enzyme with HaloMPNN raised its negatively charged surface fraction from about 20% to 35% and lowered its predicted isoelectric point from 5.7 to 4.6, shifting it substantially toward the halophilic homolog (which reaches *∼*51% negative surface residues) without fully matching it; surface positive charge and surface and core hydrophobicity decreased in parallel (Table S3).

To test HaloMPNN on an enzyme of applied interest, we redesigned the alginate lyase BoPL17 from the non-halophile *Bacteroides ovatus*, a carbohydrate-active enzyme relevant to the degradation of aquatic (algal) biomass. Redesign again increased the density of negative surface charge (Figure 2E), raising the negatively charged surface fraction from about 18% to 39% and lowering the predicted isoelectric point from 5.8 to 4.3, with accompanying decreases in surface positive charge and in surface and core hydrophobicity (Table S4).

### HaloMPNN designs proteins towards halophilic properties more effectively than SolubleMPNN, ProteinMPNN, or HyperMPNN

HaloMPNN redesign shifts the properties of proteins towards those of natural halophilic proteins, whereas SolubleMPNN, ProteinMPNN, and HyperMPNN do so only partially or not at all (Figure 3, Tables 2 and S5). HaloMPNN is the only model to move all five metrics in the halophilic direction, and it ranks the most halophilic of the four models on each metric individually (Table S5). (All differences described are significant at *q <* 10*^−^*^3^ except where stated otherwise; see Table S8.)

**Figure 3:**
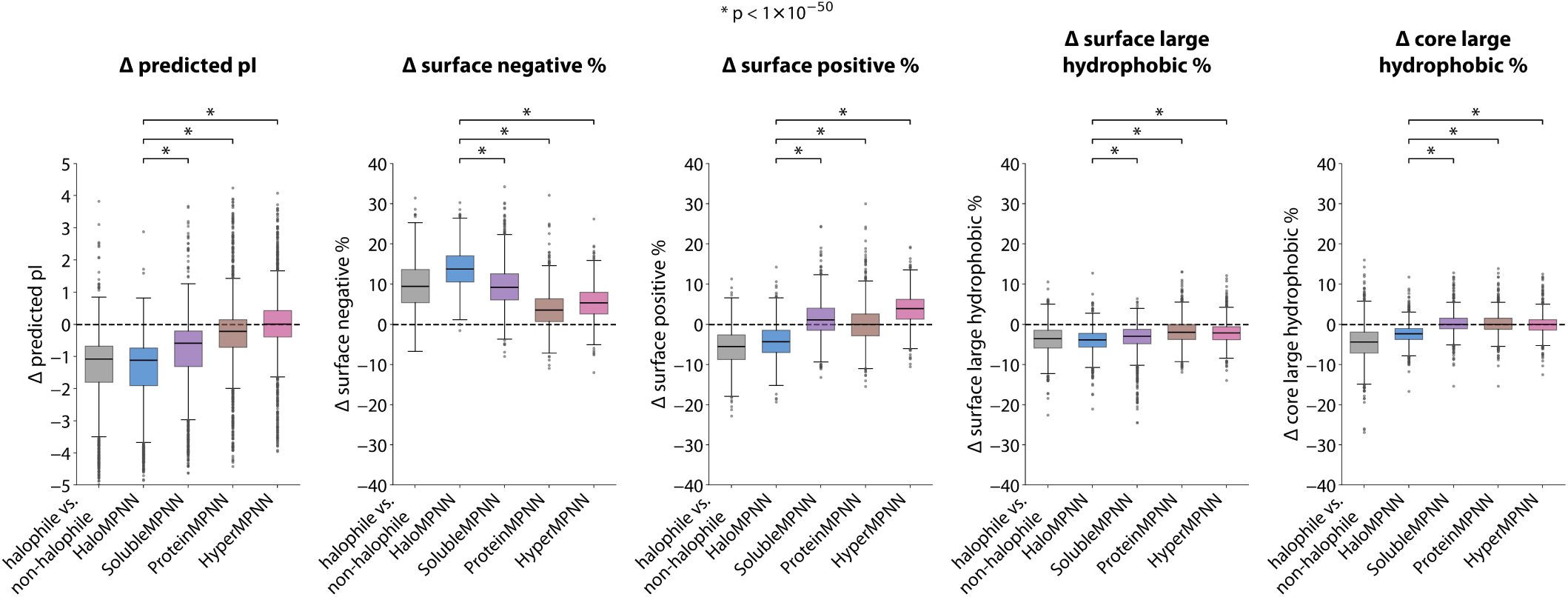
HaloMPNN designs proteins towards halophilic properties more effectively than SolubleMPNN, ProteinMPNN, or HyperMPNN. Change in predicted isoelectric point, % negatively charged surface residues, % positively charged surface residues, % large hydrophobic surface residues, and % large hydrophobic core residues from before to after redesign (N=1,433). Change in each metric between non-halophilic and halophilic homologs is shown at left for comparison. Two points were omitted from the plotted distributions only (Methods). The corresponding residue substitutions are tabulated in Tables 3, S6 and S7. Statistical test results are provided in Table S8.

HaloMPNN decreases predicted isoelectric point by an amount close to the difference between natural halophilic and non-halophilic homologs. SolubleMPNN and ProteinMPNN lower it less, and HyperMPNN leaves it essentially unchanged (*q* = 0.56).

The surface-charge metrics reveal the clearest distinction between the models. Every model increases the fraction of negatively charged surface residues, which indicates that surface acidification is a generic tendency of redesign rather than a halophilic-specific one; HaloMPNN simply drives it furthest, exceeding even the shift seen between natural homologs. The models diverge, however, on positive charge. Only HaloMPNN reduces the fraction of positively charged surface residues, approaching the natural shift, so that its redesigns acquire a net acidic surface. SolubleMPNN and HyperMPNN instead increase surface positive %, and ProteinMPNN leaves it unchanged (*q* = 0.06); these models raise the total charge of the surface without shifting its net charge toward acidic. HyperMPNN is the extreme case, increasing both negative and positive surface residues while leaving isoelectric point unchanged, a pattern more consistent with the surface salt bridges that stabilize thermophilic proteins than with the net acidic surface characteristic of halophiles.

All models reduce the fraction of large hydrophobic residues on the surface, with HaloMPNN again producing the largest decrease. The core is more discriminating. Natural halophilic proteins substantially reduce their large hydrophobic core content, and HaloMPNN is the only model to reproduce this reduction at all: SolubleMPNN increases core hydrophobicity (*q <* 10*^−^*^2^), while ProteinMPNN and HyperMPNN leave it unchanged (*q* = 0.13 and *q* = 0.09). Because the core is buried and tightly packed against the fixed backbone, altering its composition is harder than editing the surface, which may be why HaloMPNN recovers only about half of the natural core reduction even as it matches or exceeds the natural shifts in isoelectric point, surface acidity, and surface hydrophobicity.

Taken together, these comparisons separate two components of halophilic adaptation: a net acidic surface, produced by both adding negative and removing positive charge, and a reduced hydrophobic core. Only HaloMPNN captures both. The other models reproduce at most the generic surface acidification, and HyperMPNN in fact moves the surface toward a thermophilic signature. That a single change to the training distribution reorients the model across all five metrics, including the internal core change that generic redesign leaves untouched, indicates that HaloMPNN has learned features specific to halophilic adaptation rather than amplifying the default behavior of inverse folding.

### Residue substitutions in HaloMPNN redesigns recapitulate natural halophilic substitutions

To identify which substitutions drive these compositional shifts, we tabulated residue replacements between aligned native and halophilic or redesigned sequences, over all positions and separately for surface and core positions, following Paul et al. 2008^10^ (Methods, Tables 3, S6 and S7). Because the more abundant residue inflates raw counts in one direction, we ranked pairs by the net difference between forward and backward replacements. Positions conserved at *≥*50% in each protein’s alignment were held fixed during redesign (Methods), so the redesign columns describe only the non-conserved positions the models were free to change.

On the surface, the dominant pattern in natural halophilic proteins is the replacement of a basic residue by an acidic one: K*→*E (+3,279), K*→*D (+2,414), R*→*E (+1,806), and R*→*D (+1,525) are all among the ten most frequent net substitutions (Table 3). HaloMPNN reproduces this basic-to-acidic swap (R*→*E, +2,267; K*→*E, +2,104), whereas SolubleMPNN, ProteinMPNN, and HyperMPNN instead accumulate positive charge, showing net R*→*K (+2,456, +1,548, and +1,768) alongside substitutions of neutral residues by lysine (Q*→*K, S*→*K, A*→*K). This is the substitution-level counterpart of their failure to reduce surface positive %.

**Table 3:** Ten most frequent surface substitutions for each comparison, ranked by the net difference between forward (native *→* comparison) and backward (comparison *→* native) replacements; the net difference is given in parentheses. The first column compares each non-halophilic protein with its natural halophilic homolog; the remaining columns compare each non-halophilic protein with its redesign by the named model. Counts are pooled over all 1,433 aligned pairs (Methods).

|  | halophile vs.<br>non-halophile | HaloMPNN | SolubleMPNN | ProteinMPNN | HyperMPNN |
| --- | --- | --- | --- | --- | --- |
| 1 | K→E (3,279) | Q→E (2,326) | A→E (2,708) | D→E (1,581) | D→E (3,730) |
| 2 | K→D (2,414) | R→E (2,267) | R→E (2,529) | S→E (1,574) | A→E (2,716) |
| 3 | E→D (1,891) | K→E (2,104) | D→E (2,505) | R→K (1,548) | Q→E (2,002) |
| 4 | R→E (1,806) | S→E (1,889) | R→K (2,456) | Q→E (1,518) | R→K (1,768) |
| 5 | G→D (1,568) | A→E (1,637) | S→E (2,449) | S→A (1,316) | S→E (1,643) |
| 6 | R→D (1,525) | N→D (1,620) | Q→E (2,173) | R→A (1,275) | A→K (1,450) |
| 7 | N→D (1,437) | S→D (1,513) | L→E (1,387) | R→E (1,251) | Q→K (1,407) |
| 8 | S→D (1,424) | V→E (1,345) | T→E (1,238) | A→P (1,176) | A→P (1,025) |
| 9 | K→R (1,349) | L→E (1,296) | Q→K (1,173) | E→P (1,050) | Q→R (1,006) |
| 10 | K→A (1,019) | T→E (1,214) | S→K (1,130) | Q→K (948) | T→E (998) |

Beyond this gross charge swap, natural halophilic proteins show finer preferences within each charge class, and HaloMPNN reproduces their direction while the other models reverse it. Among acidic residues, halophilic proteins favour aspartate over glutamate on the surface (a net E*→*D flux; net D*→*E = *−*1,891), as does HaloMPNN (*−*609), whereas SolubleMPNN, ProteinMPNN, and HyperMPNN favour glutamate (net D*→*E of +2,505, +1,581, and +3,730). Among basic residues, halophilic proteins and HaloMPNN favour arginine over lysine at both surface (net K*→*R +1,349 and +785) and core (net K*→*R +644 and +723) positions, while the other three models show the opposite. That HaloMPNN recovers these subtler D-over-E and R-over-K preferences, and not merely the coarse addition of acidic residues, suggests it has learned halophile-specific sequence preferences rather than a generic acidification, albeit at smaller magnitude than is seen in nature.

Substitution of polar residues by acidic ones is common to all models, though the halophilic bias toward aspartate is again visible (Table 3). Q*→*E is a large net substitution in halophilic proteins and in every model, and S*→*D and S*→*E appear throughout. N*→*D, by contrast, ranks among the most frequent surface substitutions only in natural halophilic proteins (+1,437) and HaloMPNN (+1,620).

In the core, the most frequent net substitution in both natural halophilic proteins (+5,068) and HaloMPNN redesigns (+2,625) is I*→*V, a reduction in hydrophobic side-chain volume, accompanied by I*→*L (+2,407 and +922) (Table S7). This substitution is far weaker in ProteinMPNN (+404) and absent from the leading substitutions of SolubleMPNN and HyperMPNN, consistent with HaloMPNN being the only model to reduce core hydrophobic content.

### HaloMPNN generalizes to the non-halophilic *E. coli* proteome

To confirm that HaloMPNN’s redesign preferences are not specific to the paired-homolog set, we applied the same redesign and analysis to the entire non-halophilic *Escherichia coli* proteome (4,229 proteins), redesigned with HaloMPNN, SolubleMPNN, ProteinMPNN, and HyperMPNN (Methods, Figure 4, Table S9). HaloMPNN redesign moved the whole proteome toward the halophilic signature: the predicted isoelectric point of nearly every protein decreased, collapsing the natively broad, bimodal distribution into a narrow acidic peak near pH 4.5, while the fraction of negatively charged surface residues rose and the fractions of positively charged and large hydrophobic surface residues, along with large hydrophobic core residues, fell (Figure 4, Table S9).

**Figure 4:**
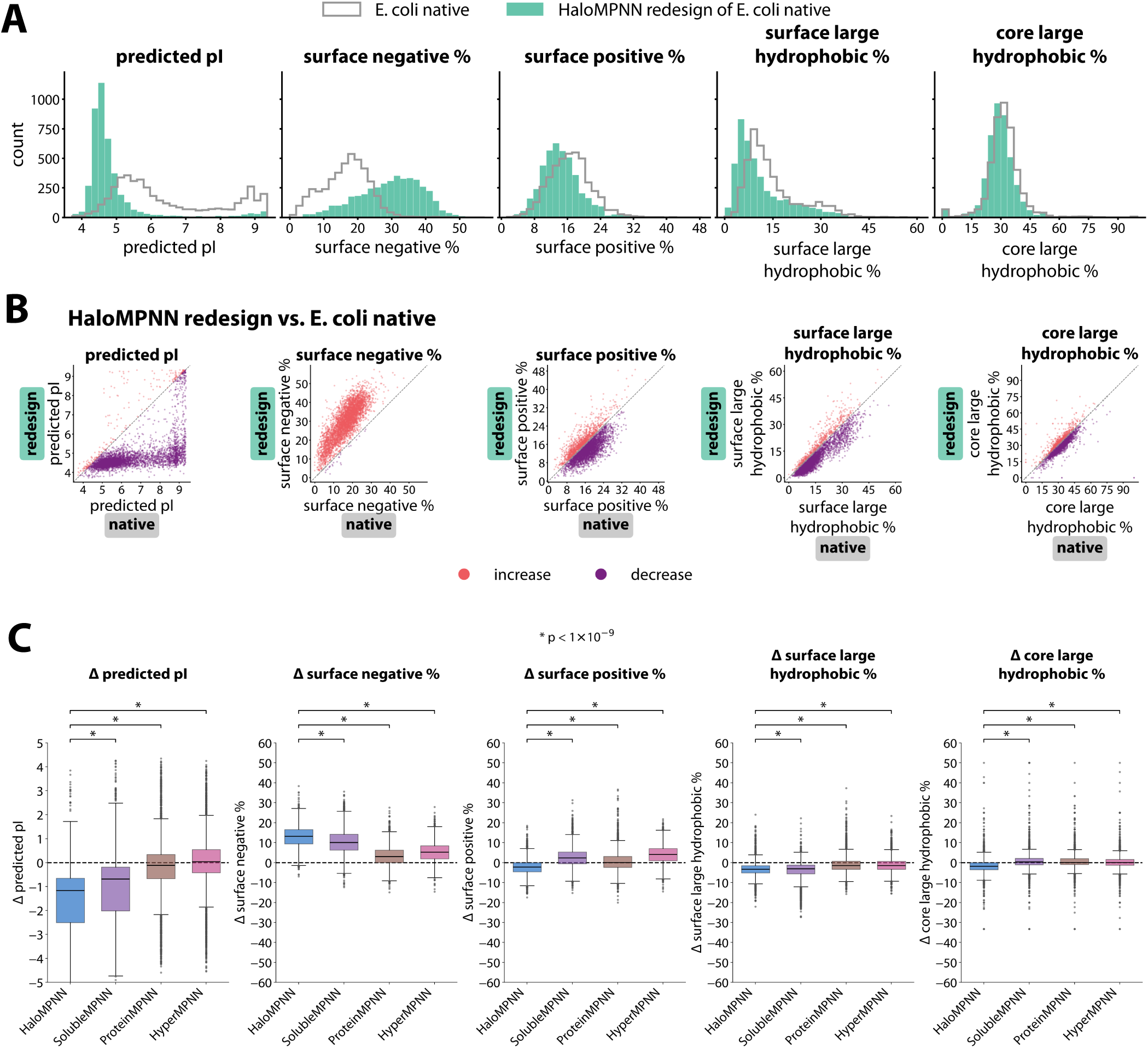
HaloMPNN imposes a halophilic signature on the non-halophilic *E. coli* proteome. Redesign of 4,229 *Escherichia coli* proteins with HaloMPNN, SolubleMPNN, ProteinMPNN, and HyperMPNN. (A,B) Distributions of predicted isoelectric point, % negatively charged surface residues, % positively charged surface residues, % large hydrophobic surface residues, and % large hydrophobic core residues before (native) and after redesign. (C) Change in each metric from before to after redesign. HaloMPNN shifts every metric toward the halophilic signature more strongly than SolubleMPNN, ProteinMPNN, or HyperMPNN.

The ranking of the models recapitulated the paired-homolog result on this independent, unpaired proteome (Figure 4, Table S9). HaloMPNN reproduced the natural halophile-to-non-halophile shift in isoelectric point most closely and produced the largest increase in surface negative charge, and, as in the paired analysis, it was the only model to decrease surface positive charge, whereas SolubleMPNN, ProteinMPNN, and HyperMPNN acidified the surface less and left positive charge unchanged or slightly increased. The residue substitutions underlying these shifts matched those seen in the paired-homolog analysis (Methods). Because the *E. coli* proteome is large, taxonomically distant from the training halophiles, and not paired to any protein used to characterize the halophilic signature, this consistency indicates that HaloMPNN imposes a halophilic signature on generic, unrelated protein backbones rather than exploiting features peculiar to the paired-homolog dataset.

## DISCUSSION

We set out to test whether retraining ProteinMPNN on halophilic proteomes yields a model that designs proteins with halophilic character. Using a dataset of 1,433 paired halophilic and nonhalophilic homologs, we first confirmed the molecular signature of halophilic adaptation: a lower isoelectric point, a more acidic and less basic surface, and reduced hydrophobicity at both the surface and the core. Redesigning the non-halophilic proteins with HaloMPNN reproduced this signature across all five metrics more completely than SolubleMPNN, ProteinMPNN, or HyperMPNN, which capture only the generic surface acidification common to the MPNN family of sequence redesign models. At the level of individual substitutions, HaloMPNN reproduced not only the pronounced exchange of basic for acidic surface residues but also the finer halophilic preferences for aspartate over glutamate and arginine over lysine, and reduced hydrophobic bulk in the core through I*→*V substitutions. These effects extended from the paired homologs to the computational redesign of the entire *E. coli* proteome. Together, the results show that a single change to the training distribution is sufficient to steer an inverse folding model toward the sequence features of halophilic adaptation.

Residue substitution preferences of HaloMPNN and halophiles can be rationalized: arginine distributes the charge of its guanidinium group over a larger area and forms more hydrogen bonds and salt bridges than lysine, interactions that stabilize proteins and may be especially favourable on a densely charged halophilic surface^18,19^. Similarly, the shorter carboxylate of aspartate coordinates surface cations and ordered water more tightly than glutamate, consistent with the hydration shell that keeps halophilic proteins soluble at high salt^12,20,21^. In the core, natural halophiles and HaloMPNN reduce hydrophobic side-chain volume through I*→*V substitutions, consistent with the need to weaken the hydrophobic driving force that high salt strengthens^10,13^. That HaloMPNN recovers these subtle, halophile-specific preferences, which generic redesign does not, argues that they are genuine features of the halophilic sequence signature.

As HaloMPNN was trained on halophilic sequences and evaluated by how closely its designs reproduce the compositional signature of halophilic proteins, so recovering that signature is to some degree expected rather than surprising. Furthermore, because the halophilic members of the evaluation pairs come from organisms represented in the training database, we cannot fully exclude that homologs of some evaluation proteins remained in the training set. Several observations mitigate these concerns. First, models trained on other structural datasets do not reproduce the signature, so it is specific to the halophilic training distribution rather than a generic consequence of redesign. Second, the redesign experiments are applied to the non-halophilic members of each pair, which are not part of the halophilic training distribution, and the agreement extends to fine-grained substitution preferences (D over E, R over K, and core I*→*V) that the coarse compositional metrics do not capture. Third, the same shifts appear on the unrelated *E. coli* proteome, which is not paired to any training protein. However, our charge analysis also does not consider the spacing and geometry of charged surface residues, which govern hydration-shell formation and salt bridges, and a direct comparison of HaloMPNN against simple surface supercharging on the same proteins would clarify whether learned redesign offers an advantage beyond charge magnitude alone. Such experiments could also resolve the longstanding question of whether increased surface acidity or decreased hydrophobicity is individually necessary or sufficient for salt tolerance.

More broadly, our results speak to the potential and limitations of the retrain-on-phenotype strategy for extremophilic design. Retraining or fine-tuning an inverse folding model on a phenotype-enriched dataset has now been applied to solubility (SolubleMPNN), hyperthermophily (HyperMPNN), alkaline adaptation (AlkalineMPNN), and, here, halophily. The experimental support behind these applications is uneven: soluble analogues of membrane proteins have been expressed and characterized^2^, and HyperMPNN designs have been reported to increase the thermostability of a self-assembling protein nanoparticle^3^, whereas the alkaline and halophilic cases so far rest on computational signatures^4^. Whether the strategy generalizes to arbitrary properties, and whether the compositional signatures it produces translate into function, therefore remains an open question. Answering it requires phenotypes with tractable experimental readouts that turn designs into concrete, testable predictions, and salt tolerance is well suited to this, since the solubility and activity of an enzyme can be measured directly across a range of salt concentrations.

With the change of training distribution, HaloMPNN recovers the compositional and substitution-level signatures of halophilic adaptation (surface acidification, loss of surface positive charge, and reduced hydrophobic bulk in the core) more completely than models trained for other properties. Designing enzymes that remain soluble and active at molar salt would allow biomanufacturing to run in seawater or brine, tolerate high-ionic-strength feedstocks, and resist contamination, capabilities central to the sustainable use of marine bioresources^6–8^. As a further instance of the retrain- on-phenotype strategy, HaloMPNN also supports the prospect that other extremophile, and even polyextremophile, adaptations may come within reach given sufficient sequenced genomes and predicted structures. Because salt tolerance is experimentally accessible, it offers a concrete test of how far the strategy can be pushed, and the wet-lab construction and characterization of HaloMPNN- generated designs is currently underway in our laboratories.

## MATERIALS AND METHODS

### Selection of salt-tolerant dataset

Most organisms living at extreme high salt are “salt-in,” meaning the salt concentration in the cytoplasm is also high. In contrast, some organisms living at elevated salt concentrations are “salt-out,” meaning the cells pump salt out so that the salt concentration in the cytoplasm is similar to mesohalophilic organisms^9,22^. The cytoplasmic proteins of “salt-in” organisms must be able to function at high salt concentrations, whereas the cytoplasmic proteins of “salt-out” organisms may not function at high salt concentrations. To increase the likelihood that the proteins in our training dataset are truly salt-tolerant, we selected proteins only from organisms labeled “extreme halophiles” from the HaloDom database, which are more likely to be “salt-in” than organisms living at “slight” or “moderate” salt concentrations^16^. To confirm that our dataset contains mainly salt-tolerant proteins, we used IPC2.0 to predict isoelectric point, which is strongly correlated with salt tolerance^23^. Proteins in the “extreme” halotolerant dataset have a lower isoelectric point than proteins in the “moderate” and “slight” categories (Figure S1).

We downloaded all protein sequences from organisms labeled “extreme” halotolerant in HaloDom^16^. We downloaded corresponding AlphaFold-predicted structures filtered for quality (pLDDT > 80) and length > 50^17,24^.

Predicted membrane proteins were identified using TMHMM ^25^ and excluded from training, following the rationale for SolubleMPNN, in which membrane proteins were withheld so that the model would not learn to place hydrophobic residues on solvent-exposed surfaces^2^. Proteins in the dataset used for evaluation (N=1,433) were also held out prior to training.

The extreme halotolerant dataset was randomly downsampled to 40,000 sequences. Sequences were clustered with MMseqs2^26^ at 0.20 sequence identity. Clusters were split into train, validation, and test sets (80%,10%,10%). The model was trained with an architecture identical to ProteinMPNN^1^, using the hyperparameters listed in Table S2; training and validation accuracy and perplexity over the course of training are shown in Figure S2.

### Paired homolog dataset from Paul et al. 2008

Proteomes were downloaded from UniProt for the sets of species identified in Paul et al. 2008^10^ (Table S1). Reciprocal best hits between each pair of species using MMseqs2^26^ search at 0.30 minimum similarity were identified as possible homologs. Possible homologs were aligned with MAFFT^27^ and filtered by *>*60% similarity and *≤*20% length difference. Proteins were then filtered for AlphaFoldDB^17^ data availability, for a final count of 1,433 halophile/non-halophile homolog pairs (Table S1).

### Redesign of non-halophilic homologs

Protein sequences from the paired halophilic/non-halophilic homolog dataset were filtered for predicted structure availability in AlphaFoldDB^17^ and length *≥*50. The resulting set of 1,433 proteins was redesigned with one sequence per target at temperature 0.1 with ProteinMPNN (v_48_020)^1^, SolubleMPNN (soluble_model_weights, v_48_020)^2^, HyperMPNN (v48_020_epoch300_hyper)^3^, and HaloMPNN. For each redesign, positions conserved at *≥*50% in the AlphaFoldDB multiple sequence alignment were held fixed at their native identity.

### Redesign of *E. coli* proteome

Protein sequences from the *Escherichia coli* proteome were downloaded from UniProt and filtered for predicted structure availability in AlphaFoldDB^17^ and length *≥*50. The resulting set of 4,229 proteins was redesigned with one sequence per target at temperature 0.1 with ProteinMPNN (v_48_020)^1^, SolubleMPNN (soluble_model_weights, v_48_020)^2^, HyperMPNN (v48_020_epoch300_hyper)^3^, and HaloMPNN, using the same conserved-position constraint.

### Surface and core residue classification

Residues were assigned to structural layers with the PyRosetta LayerSelector^28^ at its default settings, applied to the AlphaFold-predicted structure of each protein ^17,24^. The selector partitions residues into surface, boundary, and core layers using sidechain neighbor counts. The boundary layer is collapsed into the core. Natural halophilic homologs, which differ in length from their non-halophilic partners, were assigned layers from their own predicted structures. Redesigns were assigned layers from the pre-redesign non-halophilic predicted structure.

### Calculation of metrics

Predicted isoelectric point was computed from sequence with IPC2.0^23^. Surface negative (D, E), surface positive (K, R), and surface and core large hydrophobic (I, L, M, F) percentages were computed for each protein as a fraction of the residues assigned to surface or core.

### Statistical analysis

Differences between halophiles and non-halophiles (and for non-halophiles before and after redesign) were assessed with the two-sided paired sign test: for each protein the sign of the difference was recorded, pairs with a difference of exactly zero were discarded, and the number of positive differences was tested against a binomial distribution with *p* = 0.5 using scipy.stats.binomtest. Five metrics were tested across 11 comparison sets, comprising the natural homolog comparison, each of the four models against its native input, and all six pairwise comparisons between models, for 55 tests in total, each on 1,143–1,433 pairs after dropping ties. *p*-values were corrected across all 55 tests with the Benjamini–Hochberg procedure, and significance is reported on the corrected *q*-value (Table S8).

Similar tests were performed for *E. coli* redesigns (Table S9).

For display purposes only, two points with Δ core large hydrophobic % = +100, corresponding to the SolubleMPNN and ProteinMPNN redesigns of Q3B143, a small protein with few core residues, were omitted from the plotted distributions in Figure 3.

### Residue substitution analysis

Each non-halophilic protein was aligned with its natural halophilic homolog using MAFFT ^27^. In addition, redesigns were compared with native sequences. The N-terminal methionine was excluded from the substitution analysis. For each ordered pair of residues (*a, b*) we counted forward replacements (native *a* aligned to *b*) and backward replacements (native *b* aligned to *a*), and ranked pairs by the net difference between the two. Substitution tables were computed for the paired homolog dataset over all positions and separately for surface and core (Tables 3, S6 and S7) and, independently, for the redesigned *E. coli* proteome.

### Identifying example protein

To identify an illustrative example of a protein with distinct properties between halophilic and non-halophilic homologs, we filtered the 1,433 pairs so that the predicted isoelectric point of the non-halophilic homolog was less than 7.0, which is more typical of the distribution. We then selected the pair with the largest change in % negative surface residues, acetyl-CoA synthetase (Q88EH6/Q5UXS3).

### Redesign of carbohydrate-active enzyme

To demonstrate applicability to a carbohydrate-active enzyme, we additionally redesigned the alginate lyase BoPL17 (UniProt A0A1Y4PXR7) from *Bacteroides ovatus*, using its AlphaFold-predicted structure from AlphaFoldDB^17,24^. BoPL17 was selected as a potentially useful carbohydrate-active enzyme from a non-halophilic organism, since alginate lyases are of interest for the degradation of aquatic biomass. 16 sequences were generated with HaloMPNN at temperatures 0.1, 0.2, and 0.3, and the sequence with the most favorable global score was selected (temperature 0.1, score 0.9348, global score 1.2139, sequence recovery 0.3424).

### Electrostatic surface visualization

Surface charge diagrams were generated with APBS^29^ through its PyMOL interface, and the electrostatic potential was mapped onto the solvent-excluded surface. Structures used were the AlphaFoldpredicted structures of the natural proteins. For redesigned sequences, the sequence was threaded onto the backbone of the corresponding native structure by side-chain replacement only, taking the highest-probability rotamer at each substituted position with no repacking, backbone relaxation, or energy minimization.

## CODE AND DATA AVAILABILITY

Model weights and data used to generate figures are available at https://github.com/alyssa-lee/HaloMPNN_figures.

## ACKNOWLEDGMENTS

This research is partially supported by the Virtual Institute on Feedstocks of the Future (VIFF) under the Schmidt Sciences SaBRe project (Sargassum BioRefinery, S.D.K.), by the Rutgers Health BMIHAI Pilot Grant Program (R.M.), and by the National Institutes of Health, National Institute of General Medical Sciences, Award Number T32 GM135141. The authors acknowledge the Office of Advanced Research Computing (OARC) at Rutgers University.

## DECLARATION OF GENERATIVE AI USE

Claude Code (Anthropic) was used for code generation, plotting, and to assist in editing this manuscript. The authors directed the work, reviewed the code and its outputs, and take full responsibility for the content of this manuscript.

---

**Figure S1:**
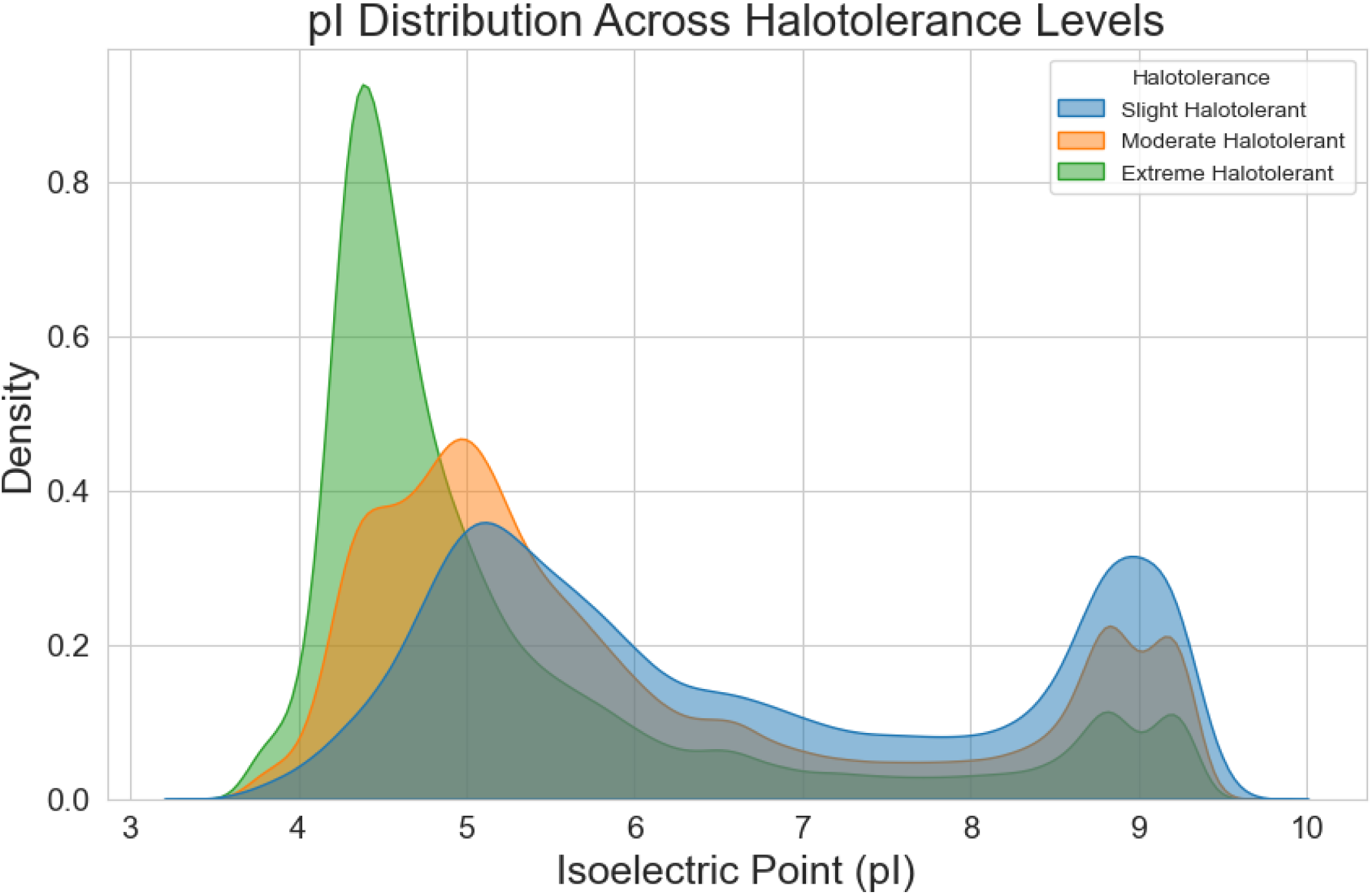
Predicted isoelectric point of proteins from HaloDom database. IPC2.0 predicted isoelectric point distributions (kernel density estimation) for proteins from HaloDom ”extreme,” ”moderate,” and ”slight” halotolerant species.

**Figure S2:**
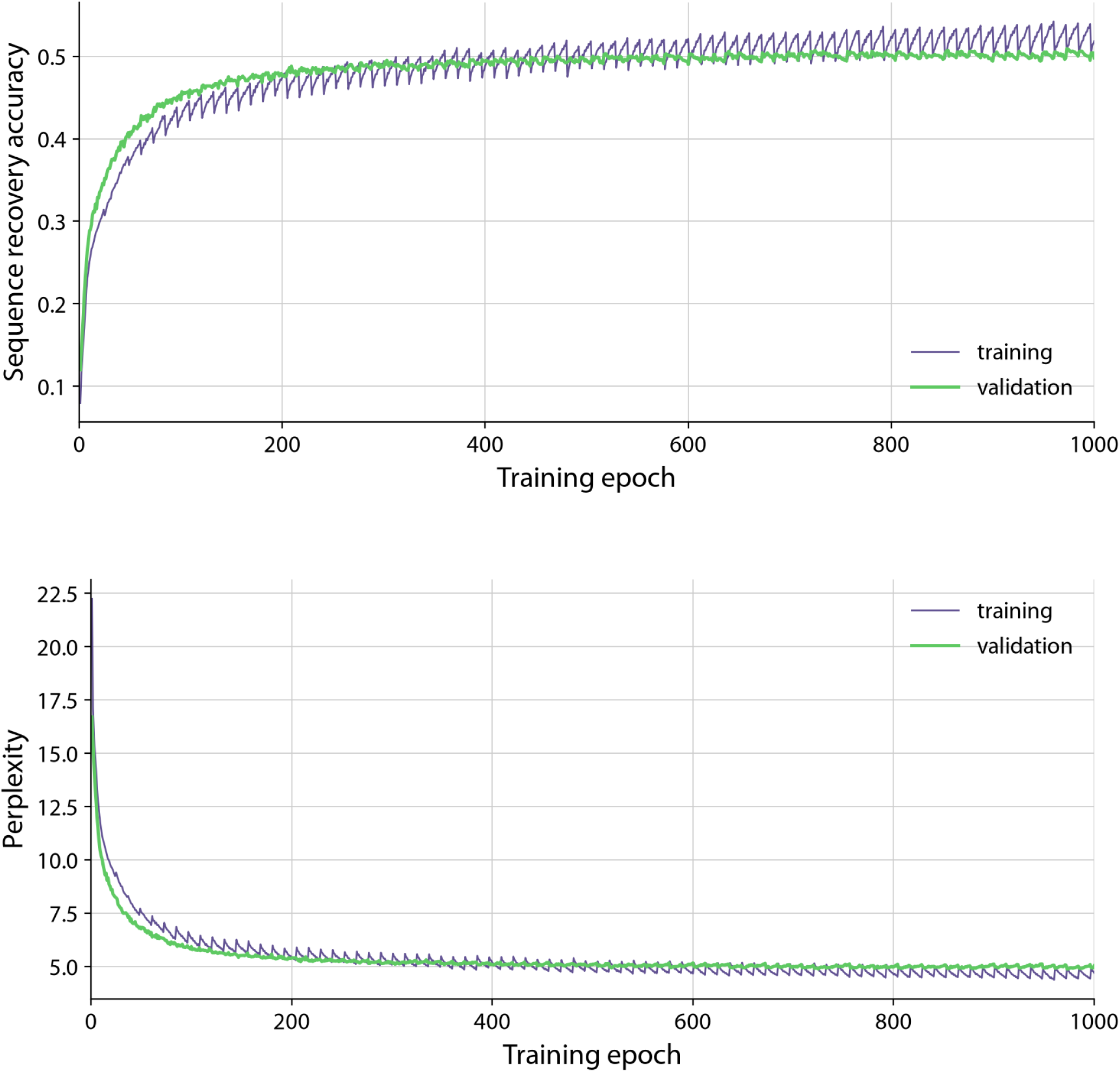
HaloMPNN training curves. Sequence recovery accuracy (top) and perplexity (bottom) on the training and validation splits over the 1,000 training epochs (57,736 optimizer steps). Final values were 0.518 (train) and 0.498 (validation) for accuracy, and 4.699 (train) and 5.028 (validation) for perplexity. Training hyperparameters are listed in Table S2.

**Table S1:**
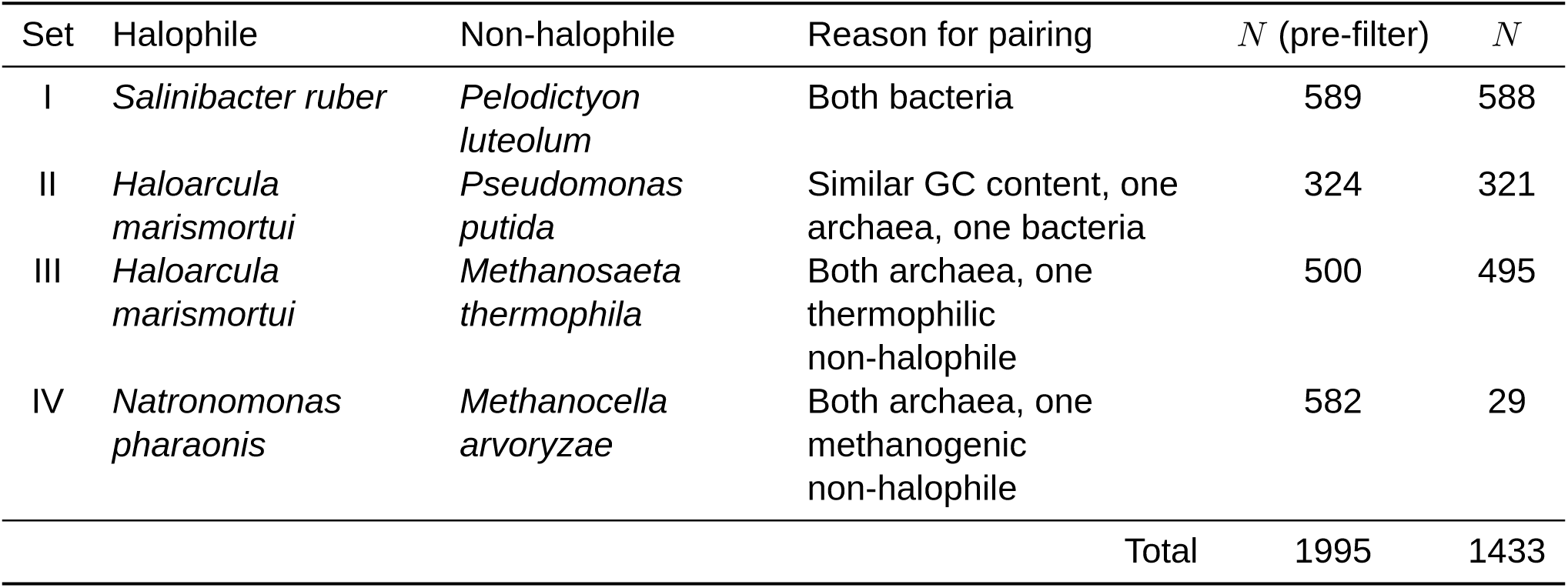
Halophile/non-halophile homolog pairs. Pairs of species chosen by Paul et al. 2008 ^10^ were reused here. Homolog pairs were computed as reciprocal best hits and filtered for AlphaFoldDB data availability (Methods).

**Table S2:** Hyperparameters used to train HaloMPNN. Parameter names follow the ProteinMPNN training script ^1^; the model architecture was left unchanged from ProteinMPNN. Training and validation curves are shown in Figure S2.

| Hyperparameter | Value |
| --- | --- |
| num_epochs | 1000 |
| save_model_every_n_epochs | 10 |
| reload_data_every_n_epochs | 2 |
| num_examples_per_epoch | 1000 |
| batch_size | 5000 |
| max_protein_length | 5000 |
| hidden_dim | 128 |
| num_encoder_layers | 3 |
| num_decoder_layers | 3 |
| num_neighbors | 48 |
| dropout | 0.1 |
| backbone_noise | 0.2 |
| rescut | 3.5 |
| gradient_norm | -1.0 |
| mixed_precision | True |

**Table S3:** Predicted pI, surface negative %, surface positive %, surface large hydrophobic %, and core large hydrophobic % for acetyl-CoA synthetase from the extreme halophile *Haloarcula marismortui* (Q5UXS3), from the non-halophile *Pseudomonas putida* (Q88EH6), and for the HaloMPNN redesign of Q88EH6. These are the proteins shown in Figure 2D.

|  | predicted pI | surface negative % | surface positive % | surface large hydrophobic % | core large hydrophobic % |
| --- | --- | --- | --- | --- | --- |
| Q5UXS3 (*H. marismortui*) | 4.05 | 51.26 | 7.54 | 4.52 | 22.15 |
| Q88EH6 (*P. putida*) | 5.69 | 19.81 | 20.29 | 6.76 | 23.54 |
| Q88EH6, HaloMPNN redesign | 4.57 | 35.27 | 14.98 | 3.86 | 22.65 |

**Table S4:** Predicted pI, surface negative %, surface positive %, surface large hydrophobic %, and core large hydrophobic % for the alginate lyase BoPL17 from the non-halophile *Bacteroides ovatus* before and after HaloMPNN redesign. These are the proteins shown in Figure 2E.

|  | predicted pI | surface negative % | surface positive % | surface large hydrophobic % | core large hydrophobic % |
| --- | --- | --- | --- | --- | --- |
| BoPL17 (*B. ovatus*) | 5.84 | 17.96 | 18.78 | 11.02 | 25.71 |
| BoPL17, HaloMPNN redesign | 4.28 | 39.18 | 10.20 | 8.16 | 22.45 |

**Table S5:** Relative ordering of values between redesigns with various MPNN models for predicted isoelectric point, surface negative %, surface positive %, surface large hydrophobic %, and core large hydrophobic %. BH-adjusted sign test was used to test significance (Table S8). * q < 0.05 ** q < 10^−2^ *** q < 10^−3^

| Metric | Ordering |  |  |  |  |  |  |
| --- | --- | --- | --- | --- | --- | --- | --- |
| predicted pI | HaloMPNN | < *** | SolubleMPNN | < *** | ProteinMPNN | < *** | HyperMPNN |
| surface negative % | HaloMPNN | > *** | SolubleMPNN | > *** | HyperMPNN | > *** | ProteinMPNN |
| surface positive % | HaloMPNN | < *** | ProteinMPNN | < *** | SolubleMPNN | < *** | HyperMPNN |
| surface large hydrophobic % | HaloMPNN | < *** | SolubleMPNN | < *** | HyperMPNN | < * | ProteinMPNN |
| core large hydrophobic % | HaloMPNN | < *** | HyperMPNN | < ** | ProteinMPNN | < *** | SolubleMPNN |

**Table S6:** Ten most frequent substitutions at all positions for each comparison, ranked by the net difference between forward (native *→* comparison) and backward (comparison *→* native) replacements; the net difference is given in parentheses. The first column compares each non-halophilic protein with its natural halophilic homolog; the remaining columns compare each non-halophilic protein with its redesign by the named model. Counts are pooled over all 1,433 aligned pairs (Methods).

|  | halophile vs. non-halophile | HaloMPNN | SolubleMPNN | ProteinMPNN | HyperMPNN |
| --- | --- | --- | --- | --- | --- |
| 1 | I→V (5,988) | I→V (3,253) | R→K (3,043) | S→A (2,566) | D→E (4,150) |
| 2 | K→E (3,820) | S→A (2,808) | A→E (2,967) | R→K (1,862) | A→E (3,064) |
| 3 | K→D (2,842) | Q→E (2,697) | D→E (2,833) | Q→E (1,794) | Q→E (2,514) |
| 4 | I→L (2,813) | R→E (2,357) | S→E (2,751) | S→E (1,779) | R→K (2,444) |
| 5 | L→V (2,352) | K→E (2,290) | R→E (2,649) | D→E (1,777) | S→E (1,849) |
| 6 | R→E (2,097) | S→E (2,096) | Q→E (2,537) | M→L (1,680) | Q→K (1,804) |
| 7 | E→D (2,056) | N→D (1,900) | M→L (1,539) | A→P (1,431) | A→K (1,761) |
| 8 | K→R (2,003) | S→D (1,755) | S→A (1,468) | R→A (1,298) | M→L (1,676) |
| 9 | G→D (1,935) | M→L (1,660) | Q→K (1,422) | R→E (1,221) | A→P (1,426) |
| 10 | S→D (1,852) | V→E (1,508) | L→E (1,380) | Q→K (1,199) | Q→R (1,377) |

**Table S7:** Ten most frequent core substitutions for each comparison, ranked by the net difference between forward (native *→* comparison) and backward (comparison *→* native) replacements; the net difference is given in parentheses. The first column compares each non-halophilic protein with its natural halophilic homolog; the remaining columns compare each non-halophilic protein with its redesign by the named model. Counts are pooled over all 1,433 aligned pairs (Methods).

|  | halophile vs.<br>non-halophile | HaloMPNN | SolubleMPNN | ProteinMPNN | HyperMPNN |
| --- | --- | --- | --- | --- | --- |
| 1 | I→V (5,068) | I→V (2,625) | M→L (1,220) | S→A (1,250) | M→L (1,158) |
| 2 | I→L (2,407) | S→A (1,761) | S→A (991) | M→L (1,197) | T→V (890) |
| 3 | L→V (1,797) | M→L (1,288) | T→V (636) | C→A (578) | F→Y (844) |
| 4 | S→A (1,143) | I→L (922) | I→L (608) | T→V (527) | R→K (676) |
| 5 | L→A (923) | C→A (797) | R→K (587) | F→Y (446) | H→Y (625) |
| 6 | M→L (894) | K→R (723) | C→A (547) | C→V (407) | S→A (586) |
| 7 | I→A (867) | M→A (514) | F→Y (450) | I→V (404) | C→A (535) |
| 8 | I→T (718) | F→L (437) | C→V (408) | G→A (355) | Q→E (512) |
| 9 | L→R (667) | T→A (405) | F→L (388) | Q→L (345) | C→V (460) |
| 10 | L→T (651) | Q→E (371) | Q→E (364) | F→L (334) | A→V (433) |

**Table S8:**
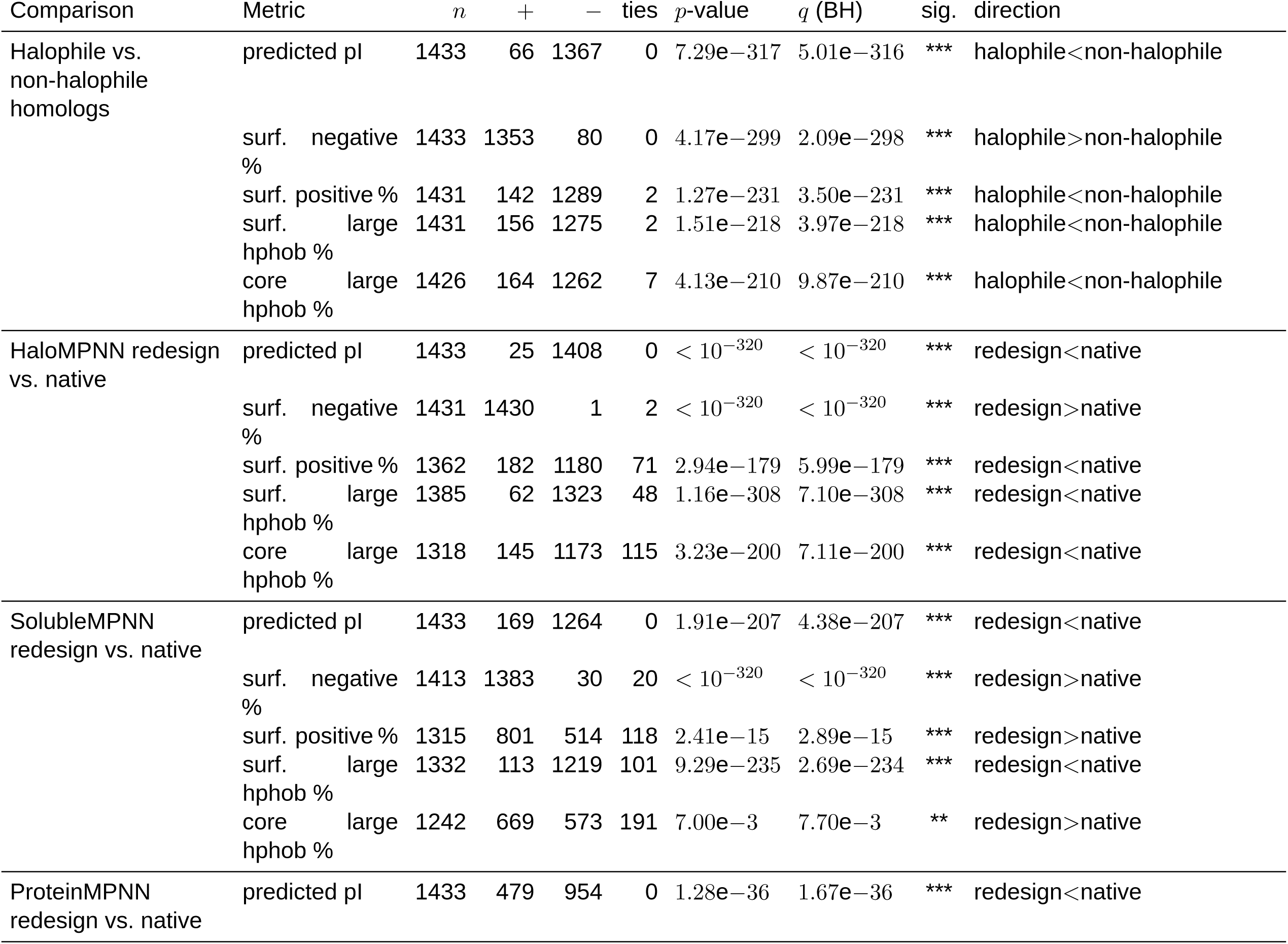
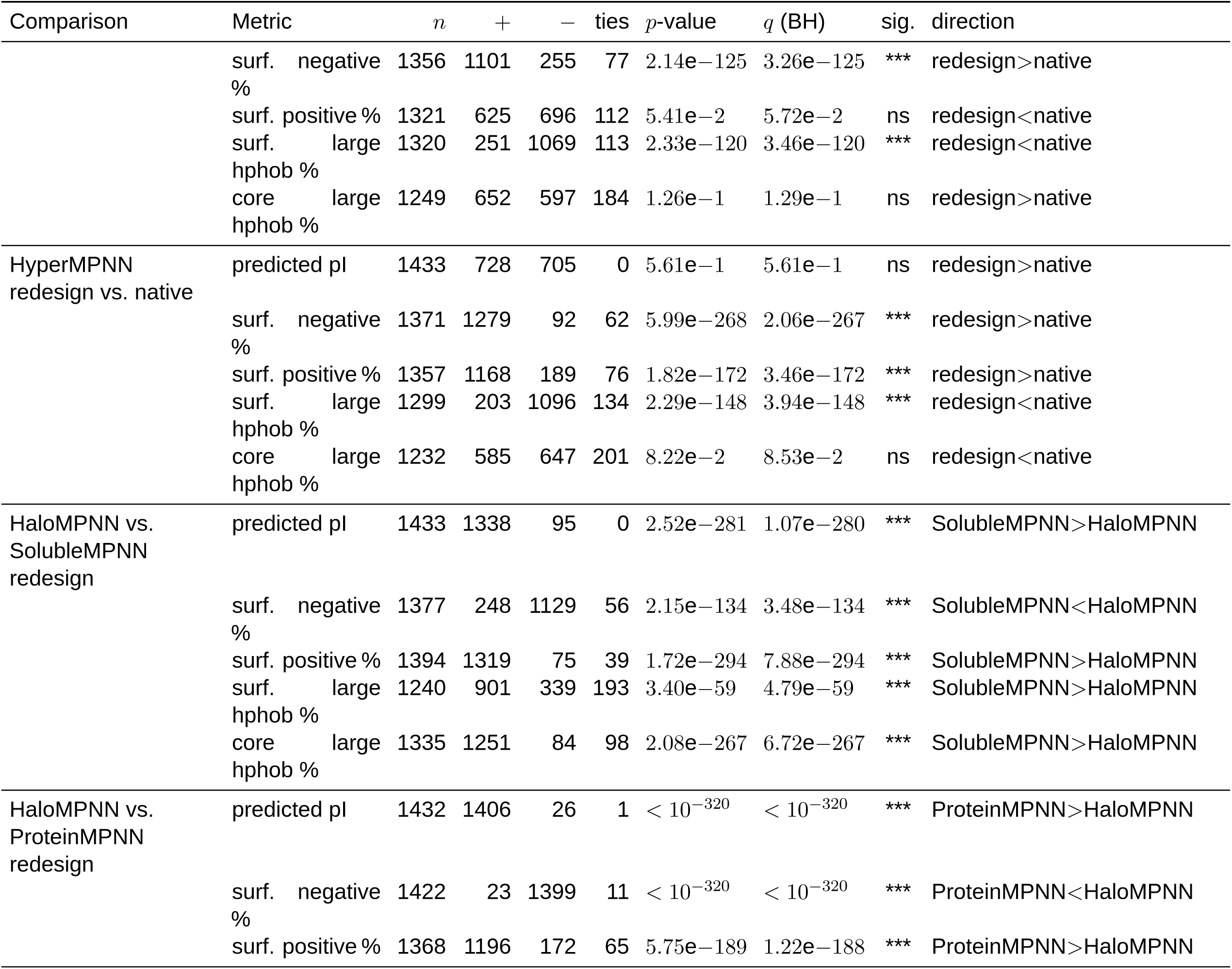
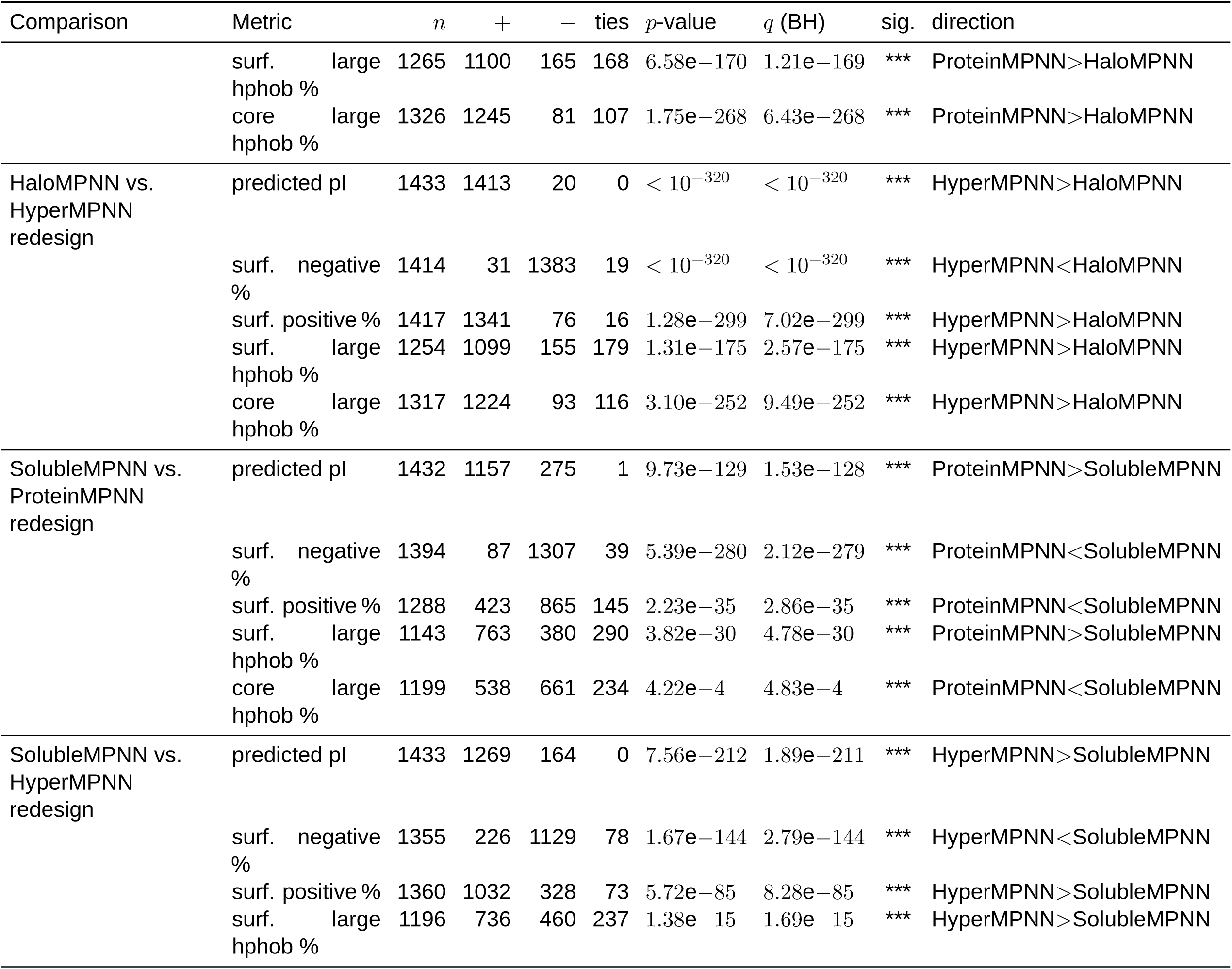
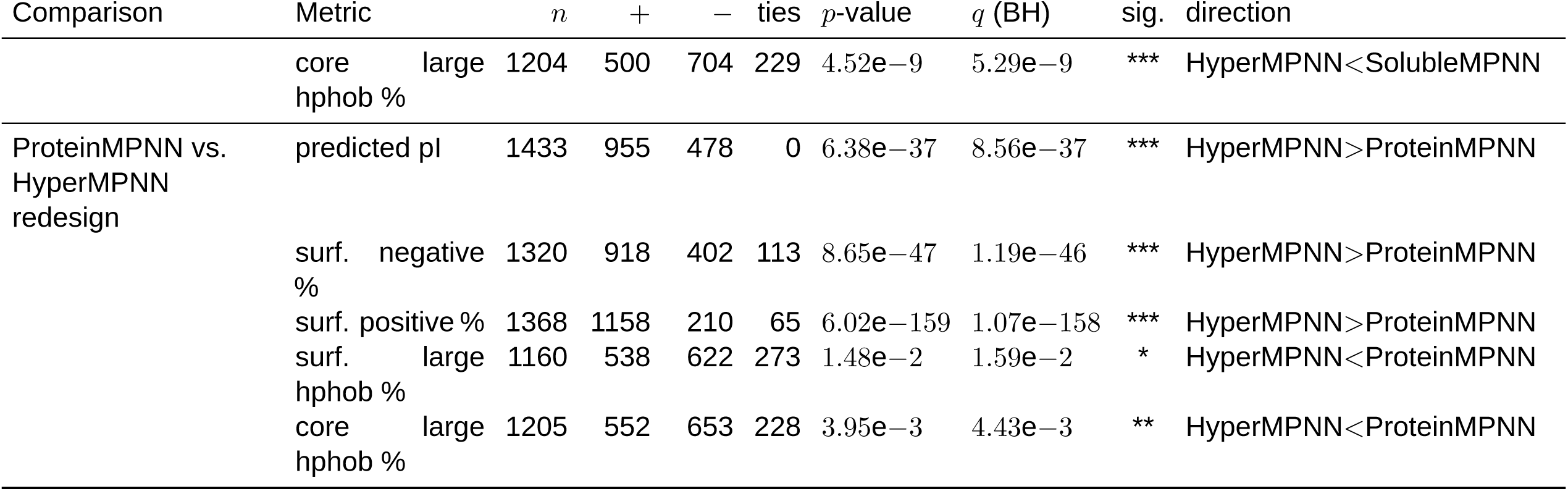
Paired sign test (two-sided binomial, ties dropped) for all metric comparisons; *q* is Benjamini–Hochberg FDR-adjusted across all tests.

**Table S9:**
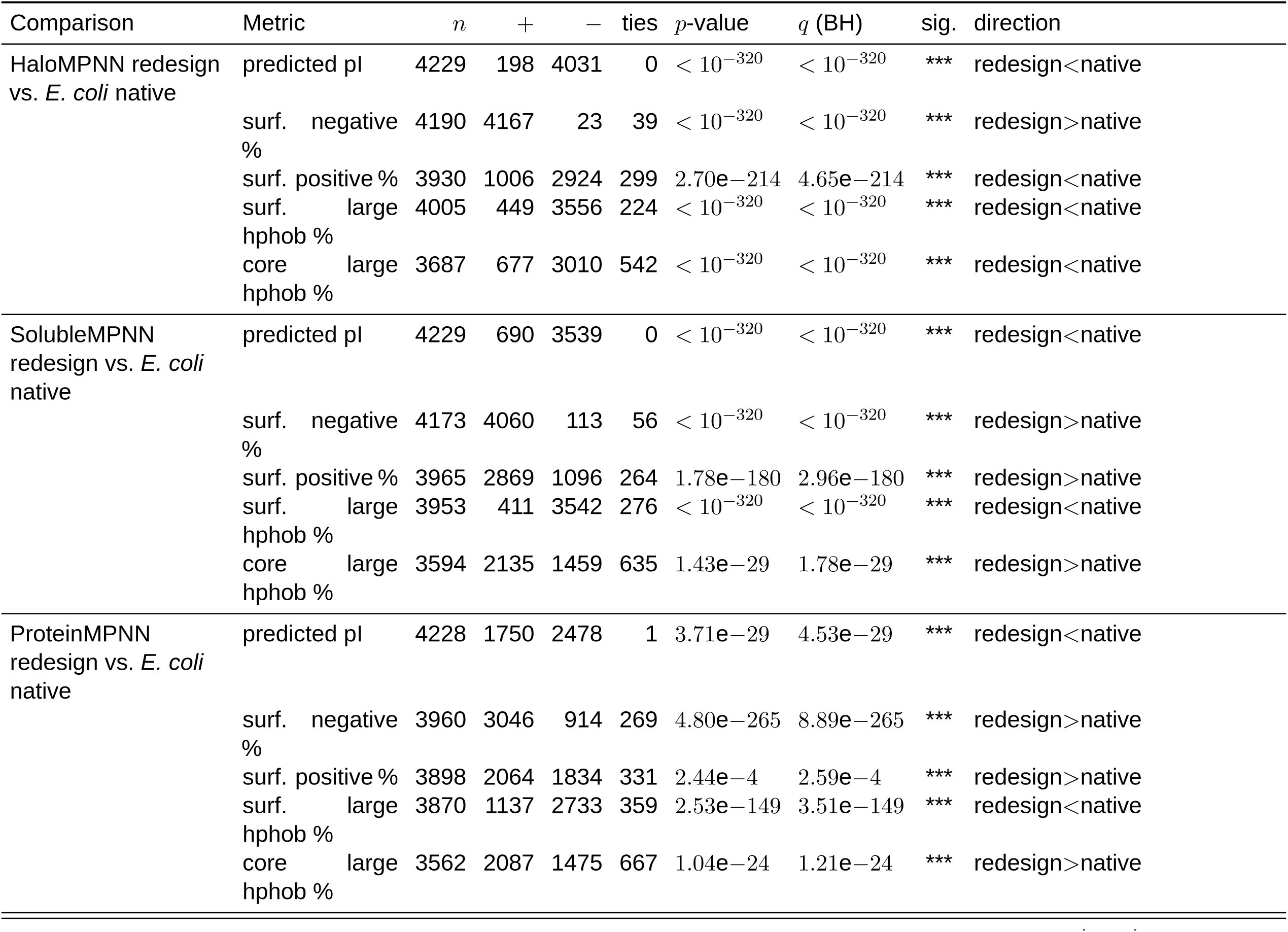
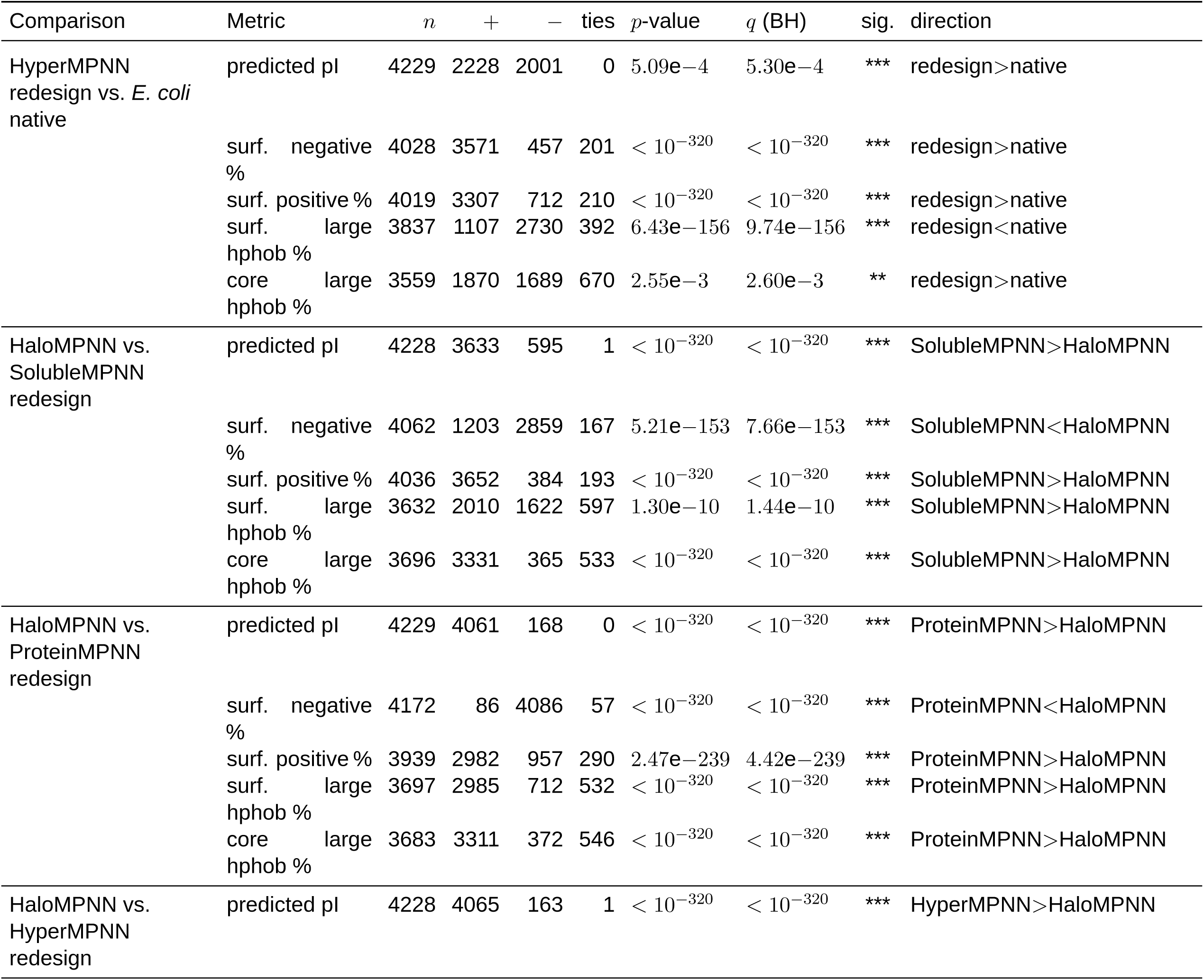
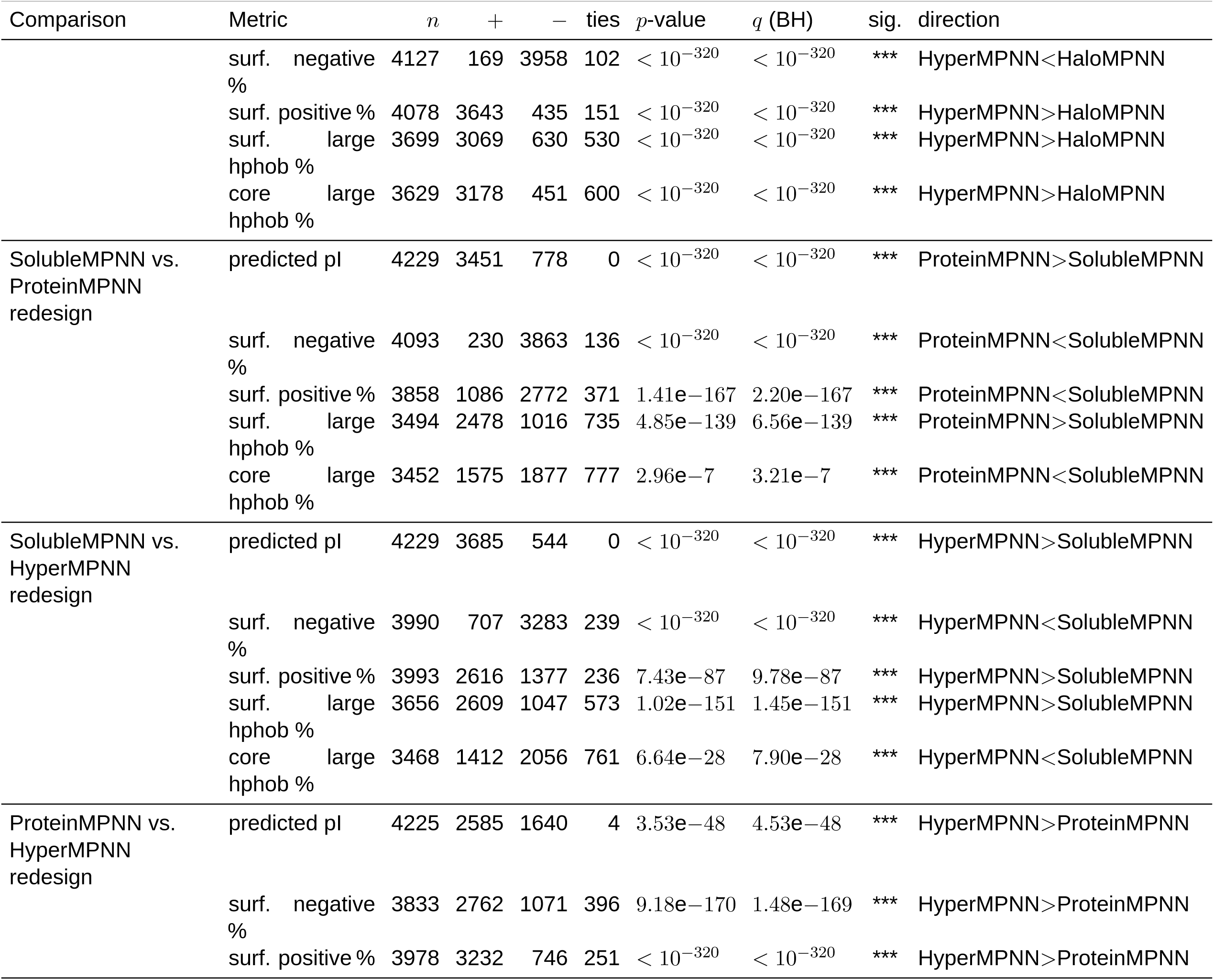
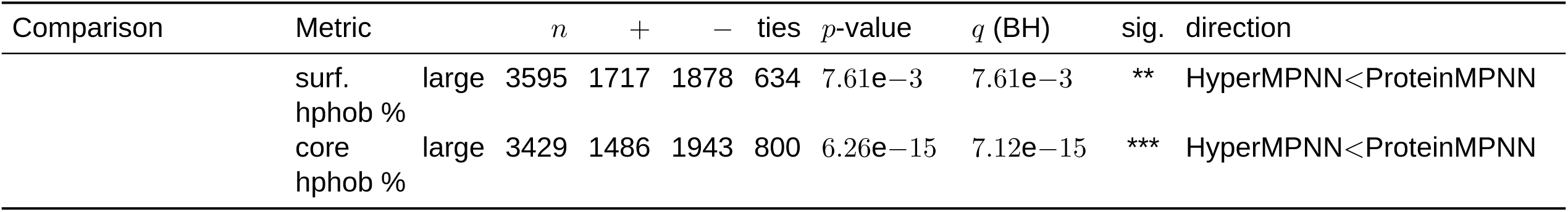
Paired sign test (two-sided binomial on sign of value2*−*value1, ties dropped) for all metric comparisons of the whole-proteome *E. coli* redesigns; *q* is Benjamini–Hochberg FDR-adjusted across all 50 tests (one global family).

